# scBasset: Sequence-based modeling of single cell ATAC-seq using convolutional neural networks

**DOI:** 10.1101/2021.09.08.459495

**Authors:** Han Yuan, David R Kelley

## Abstract

Single cell ATAC-seq (scATAC) shows great promise for studying cellular heterogeneity in epigenetic landscapes, but there remain significant challenges in the analysis of scATAC data due to the inherent high dimensionality and sparsity. Here we introduce scBasset, a sequence-based convolutional neural network method to model scATAC data. We show that by leveraging the DNA sequence information underlying accessibility peaks and the expressiveness of a neural network model, scBasset achieves state-of-the-art performance across a variety of tasks on scATAC and single cell multiome datasets, including cell type identification, scATAC profile denoising, data integration across assays, and transcription factor activity inference.

## 2 Introduction

Single cell ATAC-seq (scATAC) reveals epigenetic landscapes at single cell resolution (Buenrostro et al., 2018). The assay has been successfully applied to identify cell types and their specific regulatory elements, reveal cellular heterogeneity, map disease-associated distal elements, and reconstruct differentiation trajectories (Satpathy et al., 2019; Miao et al., 2021; Cusanovich et al., 2018).

However, there still exist significant challenges in the analysis of scATAC data, due to the inherent high dimensionality of accessible peaks and sparsity of sequencing reads per cell (Bravo Gonz’
salez-Blas et al., 2019; Chen et al., 2019). Multiple approaches have been proposed to address these challenges, which can be broadly categorized into two main classes: sequence-free and sequence-dependent methods. Starting from a sparse peak-by-cell matrix generated through aggregation of reads and peak calling in accessible chromatin, most methods represent these annotated peaks as genomic coordinates and ignore the underlying DNA sequence. Principal component analysis (PCA) and latent semantic indexing (LSI) perform a linear transformation of the peak-by-cell matrix to project the cells to a low-dimensional space (Pliner et al., 2018; Cusanovich et al., 2018). SCALE and cisTopic model the generative process of the data distribution using latent dirichlet allocation or a variational autoencoder (Bravo Gonz’
salez-Blas et al., 2019; Xiong et al., 2019). These sequence-free methods are able to detect biologically meaningful covariance to effectively represent and cluster or classify cells. However, they ignore sequence information and rely on post-hoc motif matching tools to relate accessibility to transcription factors (TFs). In contrast, sequence-dependent methods such as chromVAR and BROCKMAN represent peaks by their TF motif or k-mer content and aggregate these features across peaks or other regions of interest to learn cell representations (Schep et al., 2017; de Boer and Regev, 2018). While chromVAR directly associates peaks to TFs, emphasizing interpretability, it tends to perform worse in learning cell representations, potentially due to the loss of information from its simple implicit model relating sequence to accessibility through position weight matrices Chen et al. (2019).

Here, we propose a more expressive sequence-dependent model based on deep convolutional neural networks (CNNs). CNNs can predict peaks from bulk chromatin profiling assays more effectively than k-mer or TF motif models, exemplified by DeepSEA and Basset (Kelley et al., 2016; Zhou and Troyanskaya, 2015). These models compute explicit embeddings of the sequences underlying peaks via the convolutional layers and implicit embeddings of the multiple “tasks” (which are sequencing experiments) in parameters of the final linear transformation. We extend the Basset architecture to predict single cell chromatin accessibility from sequences, using a bottleneck layer to learn low-dimensional representations of the single cells. We show that by making use of sequence information in a deep learning framework, we outperform state-of-the-art methods for cell representation learning, single cell accessibility denoising, scATAC integration with scRNA, and transcription factor activity inference.

## 3 Results

### 3.1 scBasset predicts single cell chromatin accessibility on held-out peaks

scBasset is a deep CNN to predict chromatin accessibility from sequence. CNNs have demonstrated state-of-the-art performance for predicting epigenetic profiles in bulk data and have been successfully used for genetic variant effect prediction and TF motif grammar inference (Kelley et al., 2016; Zhou and Troyanskaya, 2015; Kelley et al., 2018; Zhou et al., 2018; Agarwal and Shendure, 2020; Avsec et al., 2021). Here, we move the focus away from maximizing accuracy on held-out sequences and view the model as a representation learning machine. When trained to achieve multiple tasks, the final layer of these models involves a sequence embedded by the convolutional layers and a linear transformation to predict the data in each separate task. The linear transfor-mation matrix comprises a vector representation of each task (here, each single cell), which specifies how to make use of each of the sequence embedding latent variables to predict cell-specific accessibility. In a simple ideal scenario, one can imagine each latent variable representing various regulatory factors such as TF binding or nucleotide composition, and the final transformation specifying how much each cell depends on that factor. We propose that these single cell vectors serve as intriguing representations of the cells for downstream tasks such as visualization and clustering.

We recommend that users first apply standard processing techniques to bring the raw data to a peak-by-cell binary matrix. scBasset takes as input a 1344 bp DNA sequence from each peak’s center and one-hot encodes it as a 4 × 1344 matrix. The input DNA sequence goes through eight convolution blocks, where each block is composed of a 1D convolution, batch normalization, max pooling, and GELU activation layers. Unlike most previous architectures, we follow these by a bottleneck layer of size *h* intended to learn a low-dimensional representation of the peak via the layer output and the cells via the parameters of the following layer. Finally, a dense linear transformation connects the bottleneck sequence embeddings to predict binary accessibility in each cell (Fig.1a). We apply the standard binary cross-entropy loss function and optimize model parameters with stochastic gradient descent (Methods).

**Figure 1:**
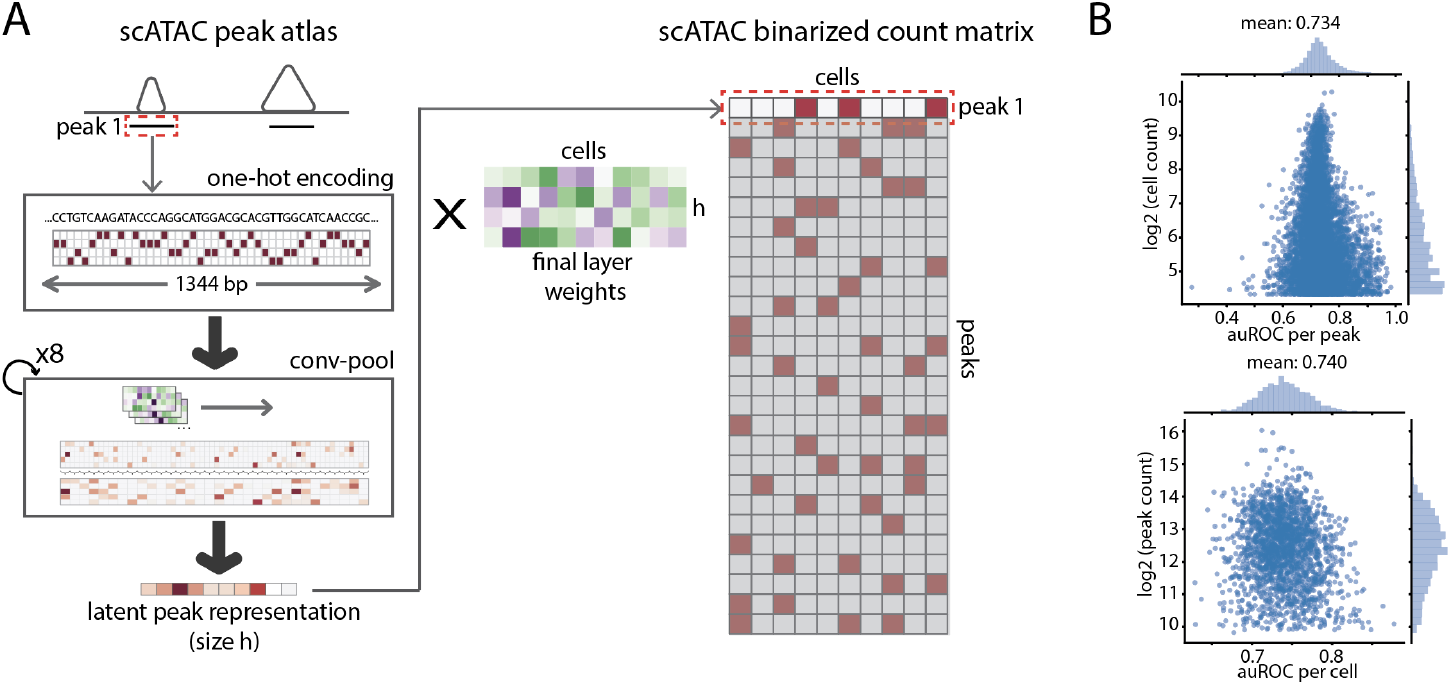
scBasset architecture. A) scBasset is a deep convolutional neural network to predict single cell chromatin accessibility from the DNA sequence underlying peak calls. B) scBasset prediction performance on held-out peaks evaluated by auROC per peak (top) and auROC per cell (bottom) for the Buenrostro2018 dataset.

To benchmark our approach, we applied scBasset to three public datasets: a scATAC-seq FACS-sorted hematopoietic differentiation dataset (referred to as Buenrostro2018) with 2k cells (Buenrostro et al., 2018), 10x Multiome RNA+ATAC PBMC dataset with 3k cells, and 10x Multiome RNA+ATAC mouse brain dataset with 5k cells. The first dataset provides ground-truth cell type labels from flow cytometry. We consider the multiome datasets to be a valuable resource to validate scATAC methods since they provide independent measurements of gene expression and chromatin accessibility in the same cells.

First, we asked how well scBasset can predict accessibility across cells for held out peak sequences to ensure that the model has learned a meaningful relationship between DNA sequence and accessibility using the sparse noisy labels. For held out peaks, we computed the area under the receiver operating characteristic curve (auROC) across peaks for each cell and averaged across cells (referred to as “per peak”). To evaluate cell type specificity, we also computed auROC across cells for each peak and averaged across peaks (referred to as “per cell”). scBasset achieved compelling accuracy levels that indicate successful learning: 0.734 per peak and 0.740 per cell for Buenrostro2018 dataset (Fig.1b), 0.662 per peak and 0.640 per cell for the 10x multiome PBMC, and 0.734 per peak and 0.701 per cell for the 10x multiome mouse brain dataset (Fig.S1). Although these statistics are slightly below the 0.75-0.95 range achieved for bulk DNase samples in the original Basset publication, this is inevitable due to the substantially increased measurement noise due to sparse sequencing for the single cell assay. In support of this claim, we observed that in the 10x multiome PBMC and mouse brain datasets, peaks with very high read coverage are easier to predict (Fig.S1). Given that ubiquitous accessible peaks are known to exist, these peaks are likely truly accessible in all cells and represent a rough upper bound on the achievable accuracy.

### 3.2 scBasset final layer learns cell representations

We propose that the *h* × cell weight matrix that connects the bottleneck layer to the predictions be used as a low-dimensional representation of the single cells. One requirement for an effective cell representation is removal of the influence of sequencing depth. Thus, we first verified that the intercept vector in the model’s final layer almost perfectly correlates with cell sequencing depth for all datasets (Fig.S2), suggesting that depth has been normalized out from the representations. Next, we compared the cell representations learned by scBasset with other methods both qualitatively and quantitatively. For the Buenrostro2018 dataset, we visualized the cell embeddings in 2D using t-distributed stochastic neighbor embedding (t-SNE) (Fig.2a) and observed differentiation trajectories in the t-SNE space. Compared to other popular methods for scATAC embedding, we observed that chromVAR and PCA have difficulty distinguishing CLP from LMPP, while Cicero, SCALE, cisTopic, and scBasset make the distinction (Fig.S4). Following previous work, we quantified the correctness of cell embeddings by comparing Louvain clustering results with ground-truth cell type labels using the adjusted rank index (ARI) (Chen et al., 2019). scBasset outperforms the other methods according to this metric (Fig.2b,top). Since ARI is sensitive to the hyperparameter choice and stochasticity in the Louvain algorithm, we proposed an alternative method for evaluating cell embeddings. We computed a “label score” by building a nearest neighbor graph based on the cell embed-dings and asked what percentage of each cell’s neighbors share its same label. For each embedding method, we computed label scores across a range of neighborhoods and observed scBasset consistently outperforms the competitors at learning cell representations that embed cells of the same type near each other (Fig.2b,bottom). We also evaluated label scores for each cell type individually and observed that monocytes are learned best, whereas MPP cells are most difficult to distinguish (Fig.S3).

**Figure 2:**
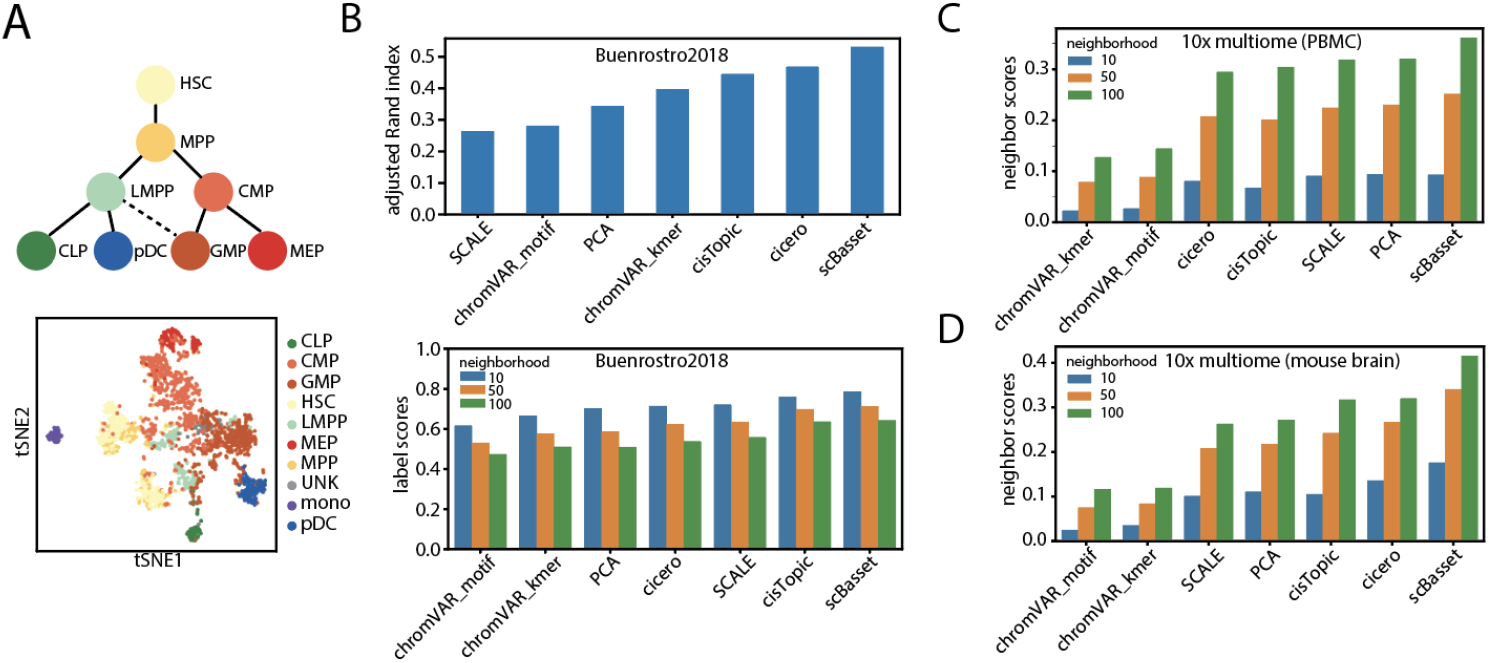
scBasset performance at learning cell representations. A) Top, hematopoietic stem cell differentiation lineage diagram in the Buenrostro2018 study; bottom, t-SNE visualization of cell embeddings learned by scBasset, colored by cell types. B) Top, performance comparison of different cell embedding methods evaluated by adjusted Rand index; bottom, performance comparison of different cell embedding methods evaluated by label score (Methods). C) Performance comparison of different cell embedding methods evaluated by neighbor scores for the 10x multiome PBMC dataset. D) Performance comparison of different cell embedding methods evaluated by neighbor scores for the 10x multiome mouse brain dataset.

For the multiome PBMC and mouse brain datasets, we computed an analogue to the label scores for cell embeddings. Since the ground-truth cell types for the multiome datasets are unknown, we used cluster identifiers from scRNA-seq Leiden clustering as cell type labels. Again, scBasset outperforms the competitors by this metric across a range of neighborhoods (Fig.S5). For these multiome datasets, we also computed a “neighbor score”, in which we built independent nearest neighbor graphs from the scRNA and scATAC and asked what percentage of each cell’s neighbors are shared between the two graphs. scBasset outperforms the competitors on both multiome PBMC and multiome mouse brain datasets when evaluated with neighbor scores across a range of neighborhoods (Fig.2c,d).

### 3.3 Batch-conditioned scBasset removes batch effects

In the Buenrostro2018 dataset, HSCs cluster into two populations, regardless of which cell embedding method we apply (Fig.S4). As noted in previous studies, this is caused by a batch effect due to different donors (Fig.3a) (Buenrostro et al., 2018; Bravo Gonz’
salez-Blas et al., 2019). To correct for this, and batch effects more generally, we explored modifications to the scBasset architecture.

**Figure 3:**
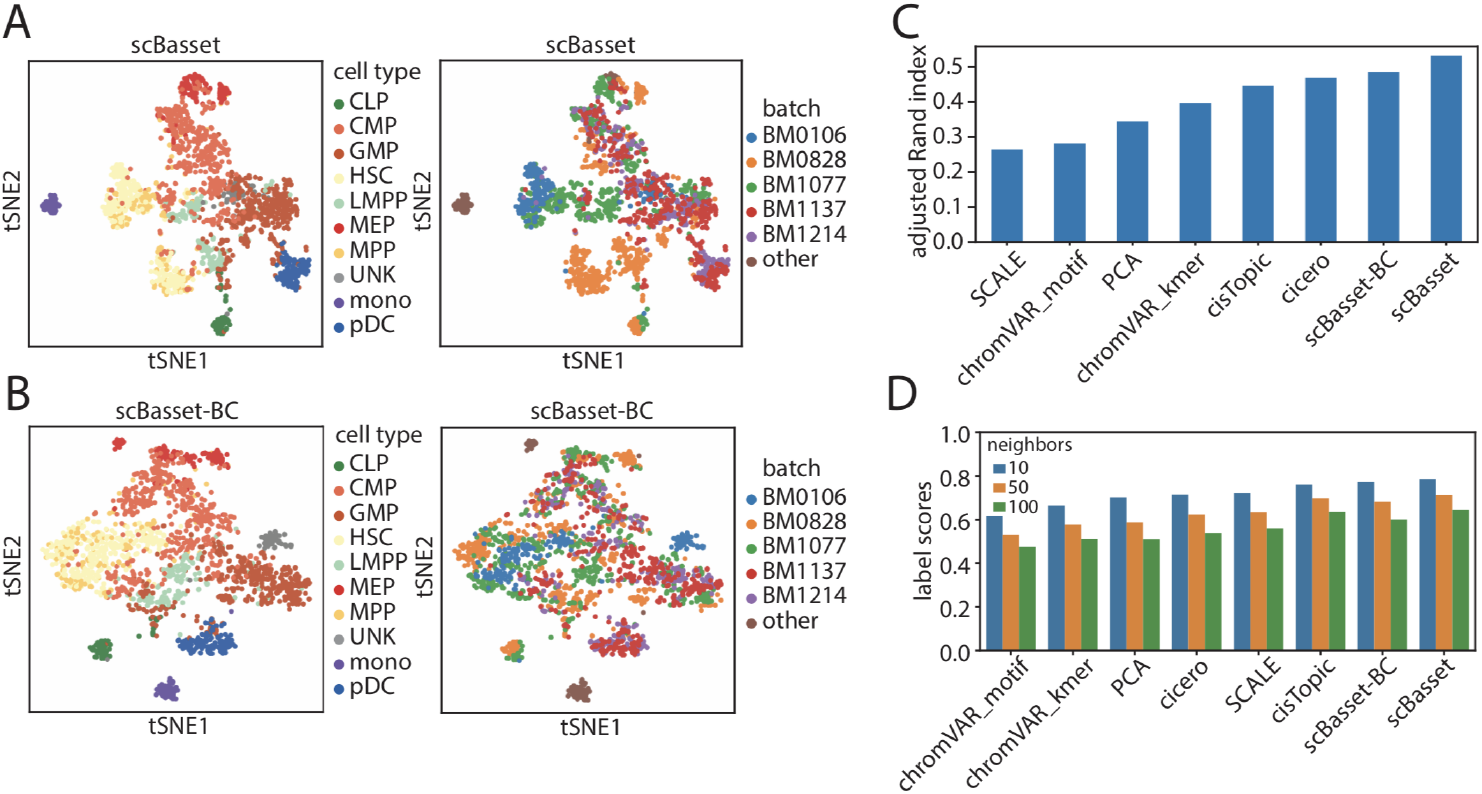
scBasset can be adapted to perform batch correction. A) Cell embeddings learned by scBasset without batch correction, colored by cell type (left) and batch (right). B) Cell embeddings learned by scBasset with batch correction (scBasset-BC), colored by cell type (left) and batch (right). C) Performance comparison of different cell embedding methods to scBasset-BC evaluated by adjusted Rand index. D) Performance comparison of different cell embedding methods to scBasset-BC evaluated by label score.

Specifically, after the bottleneck layer, we added a second fully-connected layer to predict the batch-specific contribution to accessibility (Methods, Fig.S6). We added the output of the batch layer and cell-specific layer before computing the final sigmoid. Intuitively, we expect the batch-specific variation will be captured in this path, whereas the original *h* × cell weight matrix will focus on the remainder of biologically relevant variation.

We compared the scBasset cell embedding results before and after batch correction. We observed an overall mixing of different batches in the t-SNE space after batch correction. For example, we can see that the two HSC batches (BM0106 and BM0828) merge into one cluster. In addition, pDC cells from BM1137 and BM1214 batches previously fell into two distinct sub-clusters, but are mixed together after batch correction (Fig.3ab). However, we noticed a small decrease in the cluster evaluation metrics after batch correction. We hypothesize that this is caused by imbalances in cell type distribution from different donors, which are then learned by the batch layer rather than the cell-specific layer. This is also consistent with a recent study’s observation of a trade-off between mixing and cell type separation (Ashuach et al., 2021). Nevertheless, scBasset-BC still outperforms the competitors when evaluated by ARI and is among the top performers when evaluated by label scores (Fig.3cd).

As an additional benchmark, we trained scBasset and scBasset-BC on a mixture of PBMC scATAC data from 10x multiome and 10x nextgem chemistry (Methods). We observed that while there is a strong batch effect between the two chemistries when trained with naive scBasset, scBasset-BC successfully integrated the two datasets (Fig. S6).

### 3.4 scBasset denoises single cell accessibility profiles

Due to the sparsity of scATAC, the binary accessibility indicator for any given cell and peak contains ample false negatives, such that the data cannot be studied with true single cell resolution and is usually aggregated across cells. However, numerous methods deliver denoised (or imputed) numeric values to represent the accessibility status at every cell/peak combination. scBasset computes such values in its sequence-based predictions.

From the Buenrostro2018 dataset, we sampled 500 peaks and 200 cells and directly visualized the raw cell-by-peak matrix versus the denoised matrix (Fig.4a). In the raw count matrix, we observed that cells and peaks clustered by sequencing depth, showing no biologically relevant patterns. However, we observed that after scBasset denoising, cells of the same cell type share similar accessibility profiles and hierarchical clustering of cells matched well with ground-truth labels.

**Figure 4:**
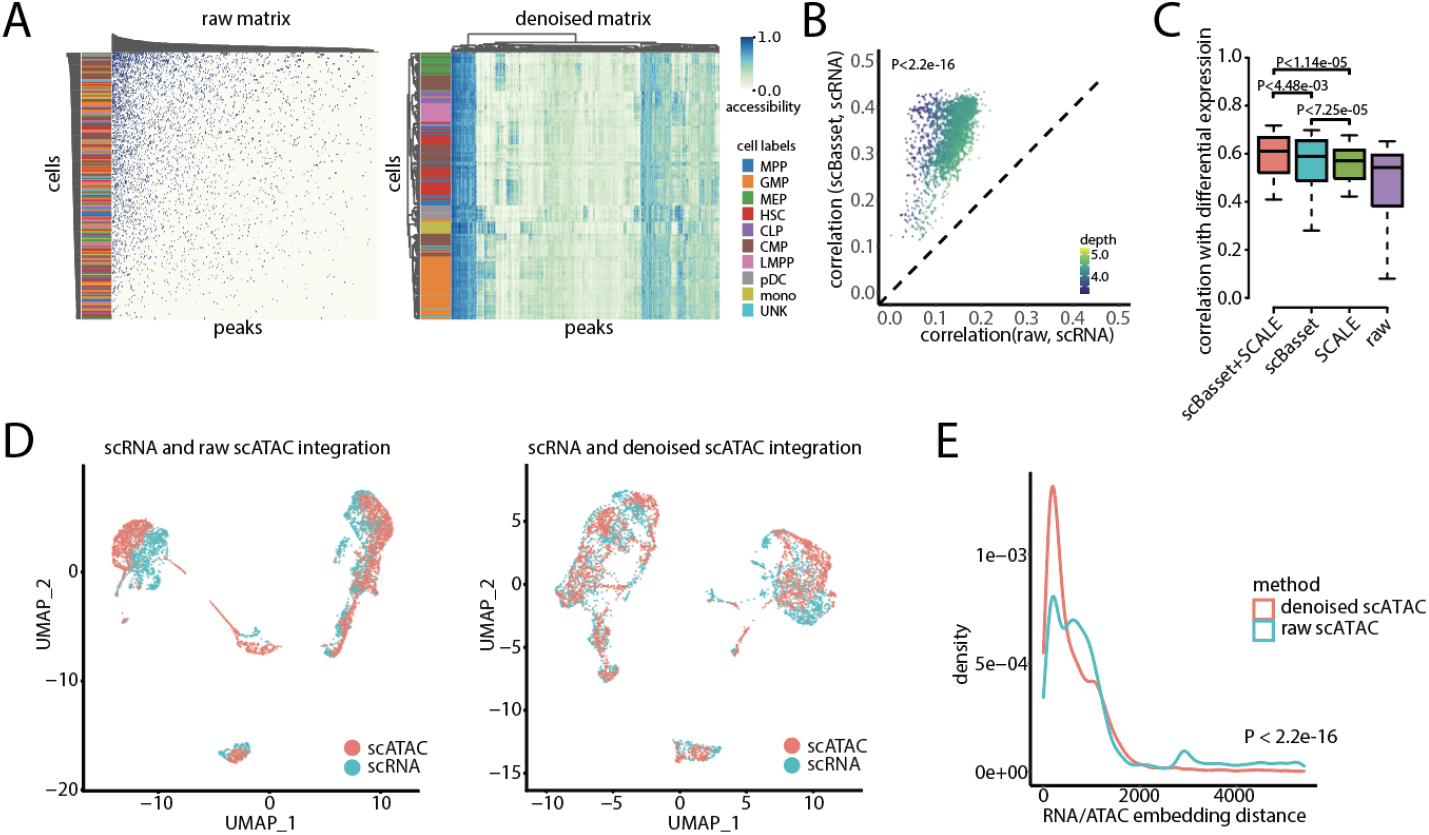
scBasset denoising performance. A) Left, binary count matrix of 200 cells and 500 peaks sampled from Buenrostro2018 dataset, hierarchically clustered by both cells and peaks. Cell type labels annotate the rows. Right, the same matrix and procedure after scBasset denoising. B) Correlation between gene accessibility score and gene expression for each cell before (x-axis) and after denoising (y-axis) for the multiome PBMC dataset. Cells are colored by sequencing depth. C) Comparison of denoising performance on multiome PBMC dataset between raw data, scBasset, SCALE, and scBasset+SCALE combine, evaluated by consistency in differential expression log2FC and differential accessibility log2FC. We performed Wilcoxon signed rank tests for performance comparisons. D) Left, 10x multiome PBMC RNA (blue) and raw ATAC (red) profile embeddings after integration. Right, 10x multiome PBMC RNA (blue) and denoised ATAC (red) profile embeddings after integration. E) Distribution of the relative distances (Method) between each cell’s RNA and ATAC embeddings after integration when using raw ATAC profiles (blue) or denoised ATAC profiles (red). We performed Wilcoxon signed rank test for performance comparison.

Several published strategies aggregate scATAC counts in the region around a gene’s transcription start site to estimate its transcription (Granja et al., 2021; Pliner et al., 2018). We propose that effective denoising would improve the correlation between these gene accessibility estimates and the gene’s measured RNA expression in multiome experiments. Thus, we computed accessibility scores for each gene by averaging the predicted accessibility values at all promoter peaks before and after denoising (Methods). For both the 10x multiome PBMC and mouse brain datasets, we observed that scBasset denoising improves the consistency between gene accessibility and expression (P<2.2e-16, Wilcoxon signed rank test). As one would expect, the improvement is greater for cells with fewer scATAC UMIs (Fig.4b, Fig.S7).

Covariance-based methods can also be used to denoise scATAC, and we compared scBasset to SCALE, a sequence-independent method for accessibility denoising using a variational autoencoder. We observed that SCALE gene accessibility scores correlated better than scBasset with gene expression (Fig.S7). Because the two methods take independent approaches (sequence-dependent versus sequence-free), we hypothesized that combining the denoised values from both via a simple average would further improve concordance. Indeed, we observed that for both 10x multiome datasets, the combined prediction performs better than SCALE or scBasset alone when we evaluated consistency with baseline expression (Fig.S7).

Studies have shown that changes in accessibility and expression correlate better with each other than their absolute values, and thus would be a more useful metric for validating accessibility denoising methods (Pliner et al., 2018). We evaluated scBasset and SCALE accessibility denoising for consistency between differential expression and differential accessibility. For each cell type cluster as defined by scRNA in the 10x PBMC dataset, we performed differential expression and differential accessibility analysis against the rest of the cells. To assess denoising quality, we evaluated the correlation between differential expression log2 fold change (log2FC) and differential accessibility log2FC before and after denoising (Fig.4c).

We observed that expression log2FC and accessibility log2FC correlates well even for raw accessibility data (r=0.47). Still, consistency is significantly improved after scBasset denoising (r=0.54). Interestingly, we observed that even though SCALE correlation exceeded that of scBasset for baseline accessibility/expression, scBasset significantly outperforms SCALE when evaluated by differential accessibility/expression (p<7.25e-05). We hypothesize that SCALE’s reliance on cell-cell covariance encourages cells to be more similar to each other than they actually are and over-smooths (Tjarnberg et al., 2021; Ashuach et al., 2021). scBasset will be less prone to over-smoothing since each peak is considered only through its sequence. As a result, SCALE performs better in denoising baseline accessibility, while scBasset performs better in denoising differential accessibility, which emphasizes cell identity. As with baseline expression, combining scBasset and SCALE produces greater performance than either method alone (Fig.4c, Fig.S7).

Integration of cells independently profiled by scRNA and scATAC into a shared latent space is a key step for many scATAC annotation and analysis methods (Stuart et al., 2019). We hypothesized that scATAC denoising would improve scRNA and scATAC integration performance. In order to evaluate integration performance, we treated the 10x multiome scRNA and scATAC profiles as having originated from two independent experiments. For the 10x multiome PBMC dataset, we observed that when we integrated the scRNA profiles with the denoised scATAC profiles, the cells achieve better mixing compared to when we integrated scRNA with raw scATAC profiles (Fig.4d). Quantitatively, we measured the multiome rank distance between the RNA and ATAC embeddings for each matching cell (Methods). We observed the RNA and ATAC profiles of the same cell are embedded significantly closer to each other when the ATAC profile is denoised compared to the raw ATAC profile (Fig.4e, P<2.2e-16). We observed similar results for the 10x multiome mouse brain dataset (Fig.S8).

### 3.5 scBasset infers transcription factor activity at single cell resolution

Transcription factor binding is a major driver of chromatin accessibility (Thurman et al., 2012). Since scBasset learns to predict accessibility from sequence, we expect the model to capture sequence information predictive of TF binding. To query the single cell TF activity, we leveraged the flexibility of the scBasset model to predict arbitrary sequences. More specifically, we fed synthetic DNA sequences (dinucleotide shuffled peaks) with and without a particular TF motif of interest to a trained scBasset model and evaluated the activity of the motif in each cell based on changes in predicted accessibility (Methods) (Kelley et al., 2016). If a TF is playing an activating role in a particular cell, we expect to see increased accessibility after the TF motif is inserted.

TF regulation in the hematopoeitic lineage profiled in the Buenrostro2018 dataset has been studied in detail. We performed motif injection for all 733 human CIS-BP motifs using the Buenrostro2018-trained model and recapitulated known trajectories of motif activity. For example, CEBPB, a known regulator of monocyte development, shows the highest activity in monocytes; GATA1, a key regulator of the erythroid lineage, is predicted to be most active in MEPs; HOXA9, a known master regulator of HSC differentiation, has highest predicted activity in HSCs (Fig.S9) (Buenrostro et al., 2018).

Previous sequence-based methods such as chromVAR are also able to quantify TF motif activity. To comprehensively compare scBasset and chromVAR on this task, we analyzed the 10x PBMC multiome dataset, in which TF expression measured in the RNA can serve as a proxy for its motif’s activity. We inferred motif activity for all 733 human CIS-BP motifs using both scBasset and chromVAR. For the 203 TFs that are significantly differentially expressed between cell type clusters, we asked how well the inferred TF activity per cell correlates with its expression. We observed that overall scBasset TF activities correlate significantly better with expression than chromVAR TF activities (P<3.38e-02, Wilcoxon signed rank test) (Fig.5b). This one-sided test is an underestimate of scBasset’s performance advantage over chromVAR, since we would expect TF expression and inferred activity to be negatively correlated for repressors. Thus, we evaluated scBasset and chromVAR on activating and repressive TFs separately. For 74 TFs which both methods agreed on a positive TF expression-activity correlation, scBasset predicted TF activities have significantly greater correlation with expression than chromVAR predicted activity (P<7.38e-12, Wilcoxon signed rank test, Fig.S10). For 41 TFs which both methods agreed on a negative TF expression-activity correlation, scBasset predicted TF activities have a significantly lesser correlation (more negative) with expression than chromVAR predicted activity (P<1.62e-08, Wilcoxon signed rank test). This is also true for the 10x multiome mouse brain dataset (Fig.S10).

**Figure 5:**
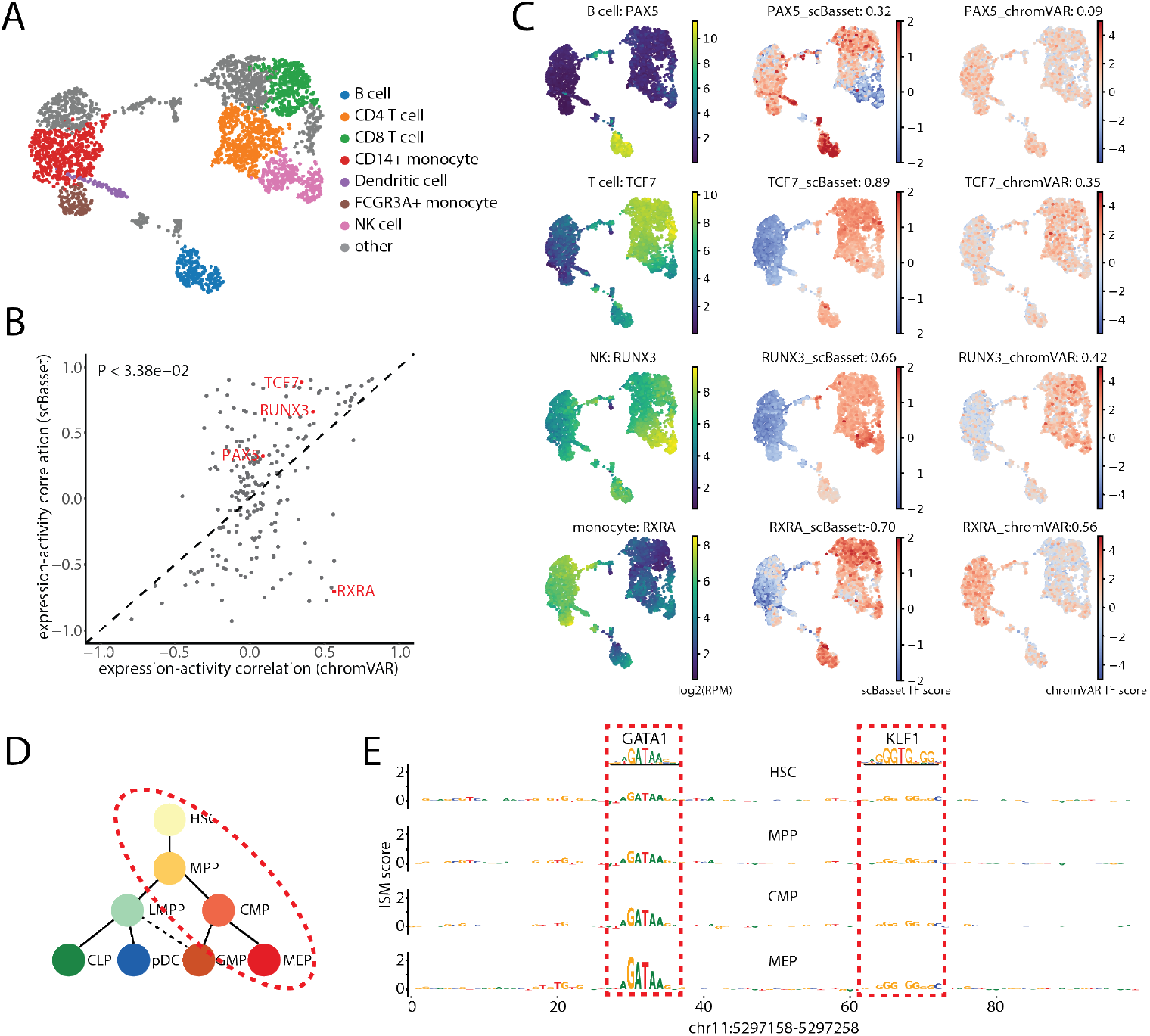
scBasset infers single cell TF activity. A) UMAP showing annotated PBMC cell types. B) Pearson correlation between TF expression and scBasset or chromVAR-predicted TF activity for 203 differentially expressed TFs. The example TFs that we examine in panel C are highlighted in red. C) UMAP visualization of TF expression (left), scBasset TF activity (middle), and chrom-VAR TF activity (right) for key PBMC regulators. Pearson correlation between inferred TF activity and expression are shown in the title. D) ISM scores for *β*-globin enhancer at chr11:5297158-5297258 for cells in HSC, MPP, CMP and MEP cell types. Sequences that match GATA1 and KLF1 motifs are highlighted in red boxes

Examining some of the key regulators of PBMC cell types, we observed that scBasset TF activities have better cell type specificity and correlate better with TF expression than chromVAR (Fig.5c). For example, PAX5 is a known master regulator of B cell development(Medvedovic et al., 2011). scBasset predicts B cell specific activity of PAX5, which correlates with PAX5 expression (r=0.32), while chromVAR PAX5 activity did not have any cell type specificity or significant PAX5 expression correlation (r=0.09). scBasset-predicted activity of the T cell differentiation regulator TCF7 highly correlates with expression (r=0.89), while chromVAR TCF7 activity has lesser specificity and expression correlation (r=0.35). NK cells have greater expression of RUNX3 and scBasset captures this elevated activity in NK cells (r=0.66) more effectively than chromVAR (r=0.42). For monocytes, both scBasset and chromVAR predicted specific activity of CEBPB, with scBasset activity correlating slightly better with expression (0.75 vs. 0.68, Fig.S11). Interestingly, while scRNA-seq suggests monocyte-specific expression of RXRA, scBasset and chromVAR strongly disagree, making opposite predictions for RXRA activity; scBasset predicts RXRA as a repressor (r=-0.70) while chromVAR suggests an activating role (r=0.56). A literature review revealed stronger evidence that RXRA plays a repressive role in the myeloid lineage through direct DNA binding, which is more consistent with the scBasset prediction (Kiss et al., 2017).

Unlike chromVAR, scBasset makes use of an accurate quantitative model that predicts cell type specific accessibility from the DNA nucleotides. Not only are we able to query scBasset for TF activity on a per-cell level, we can also infer TF activity at per-cell per-nucleotide resolution. As a proof of principle, we examined a known enhancer for the *β*-globin gene that regulates erythoid-specific *beta*-globin expression (Tuan et al., 1985; Li et al., 2002). We performed *in silico* saturation mutagenesis (ISM) for this 100 bp sequence, in which we predicted the change in accessibility in each cell after mutating each position to its three alternative nucleotides. We aggregated to a single score for each position by taking the normalized ISM score for each reference nucleotide (Methods). Fig.5d shows the average ISM score for each cell type in the erythroid lineage. We observed that the most influential nucleotides correspond to GATA1 and KLF1 motifs, which are TFs known to bind to this enhancer region and regulate *β*-globin expression (Tallack et al., 2010). Interestingly, GATA1 and KLF1 motifs contribute more to the accessibility as the cells differentiate in the erythroid lineage. In comparison, these two motifs’ nucleotides have low scores in cell types outside of the erythroid lineage (Fig.S12). This experiment suggests that scBasset learns the accessibility regulatory grammar at single cell resolution and could be used to identify the TFs regulating specific enhancers in individual cells and lineages.

## 4 Discussion

In this study we present scBasset, a sequence-based deep learning framework for modeling scATAC data. scBasset is trained to predict individual cell accessibility from the DNA sequence underlying ATAC peaks, learning a vector embedding to represent the single cells in the process. A trained scBasset model can strengthen multiple lines of scATAC analysis, and we demonstrate state-of-the-art performance on several tasks. Clustering the model’s cell embeddings achieves greater alignment with ground-truth cell type labels. The model outputs can be used as denoised accessibility profiles, which improve concordance with RNA measurements. The model learns to recognize TF motifs and their influence on accessibility, and we designed an *in silico* experiment to inject motifs into background sequences to query for TF motif activity in single cells. The model can also be applied to predict the influence of mutations, enabling *in silico* saturation mutagenesis of regulatory sequences of interest at single cell resolution. Compared to previous sequence-based approaches for scATAC analysis such as chromVAR, scBasset achieves better performance at learning cell embeddings and inferring TF activity, because scBasset benefits from a more expressive CNN model that learns more sophisticated sequence features, including non-linear relationships. Compared to previous sequence-free approaches such as cisTopic or SCALE, scBasset achieves better performance in benchmarking tasks and delivers a more interpretable model that can be directly queried for TF activity or identifying regulatory sequences.

Sequence-based approaches have several limitations. First, we make use of the reference genome, but many samples will have variant versions, including copy number variations that could lead our models astray. Second, we assume that the regulatory motifs and their interactions generalize across the genome. This assumption may not be entirely true at some genomic loci for which evolution led to bespoke regulatory solutions, such as for X chromosome inactivation in females. However, since scBasset takes a completely independent approach to the covariance-based methods, one can always combine these two types of approaches to further improve their analyses, as we showed for denoising (Fig.4).

In addition, we foresee several paths to further improve our method. To improve scBasset memory efficiency in order to scale to extremely large datasets, one could sample mini-batches of both sequences and cells rather than only sequences in our current implementation. Methods such as TF-MoDISco could be applied to scBasset ISM scores for de novo motif discovery (Shrikumar et al., 2018; Avsec et al., 2021). All approaches to scATAC analysis depend on accurate peak calls, and predictive modeling frameworks have been proposed to help identify highly specific regulatory elements (Lal et al., 2021). We expect a neural network model would further improve scATAC peak calling by taking into account sequence information (and accounting for Tn5 transposition bias). Finally, we plan to explore transfer learning approaches in which models are pre-trained on large data compendia before fine-tune training on specific single cell datasets.

## 5 Methods

### 5.1 scATAC-seq preprocessing

We downloaded the count matrix and peak atlas files for the Buenrostro2018 dataset from GEO (Accession GSE96769) (Buenrostro et al., 2018). Peaks accessible in less than 1% cells were filtered out. The final dataset contains 126,719 peaks and 2,034 cells.

We downloaded the 10x multiome datasets from 10x Genomics: https://support.10xgenomics.com/single-cell-multiome-atac-gex/datasets/2.0.0/pbmc_granulocyte_sorted_3k for PBMC dataset, and https://support.10xgenomics.com/single-cell-multiome-atac-gex/datasets/2.0.0/e18_mouse_brain_fresh_5k for mouse brain dataset. Genes expressed in less than 5% cells were filtered out. Peaks accessible in less than 5% cells were filtered out.

### 5.2 scRNA-seq preprocessing

For the 10x multiome datasets, we processed the expression data with scVI version 0.6.5 with n layers=1, n hidden=768, latent=64 and a dropout rate of 0.2 (Lopez et al., 2018). We trained scVI for 1000 epochs with learning rate of 0.001, using the option to reduce the learning rate upon plateau using options lr patience of 20 and lr factor of 0.1. We enabled early stopping when there was no improvement on the ELBO loss for 40 epochs.

To generate denoised expression profiles, we used the get sample scale() function to sample from the generative model 10 times and took the average. We used the learned latent cell representations to build nearest neighbor graphs and perform cell clustering.

### 5.3 Model architecture

scBasset is a neural network architecture that predicts binary accessibility vectors for each peak based on its DNA sequence. scBasset takes as input a 1344 bp DNA sequence from each peak’s center and one-hot encodes it as a 1344 × 4 matrix. The neural network architecture includes the following blocks:

- 1D convolution layer with 288 filters of size 17 × 4, followed by batch normalization, Gaussian error linear unit (GELU), and width 3 max pooling layers, which generates a 488*×*288 output matrix.
- Convolution tower of 6 convolution blocks each consisting of convolution, batch normalization, max pooling, and GELU layers. The convolution layers have increasing numbers of filters (288, 323, 363, 407, 456, 512) and kernel width 5. The output of the convolution tower is a 7*×*512 matrix.
- 1D convolution layer with 256 filters with kernel width 1, followed by batch normalization and GELU, The output is a 7*×*256 matrix, which is then flattened into a 1*×*1792 vector.
- Dense bottleneck layer with 32 units, followed by batch normalization, dropout (rate=0.2), and GELU. The output is a compact peak representation vector of size 1*×*32.
- Final dense layer predicting continuous accessibility logits for the peaks in every cell.
- (Optional) To perform batch correction, we attach a second parallel dense layer to the bottleneck layer predicting batch-specific accessibility. This batch-specific accessibility is multiplied by the batch-by-cell matrix to compute the batch contribution to accessibility in every cell. This vector is then added to the previous continuous accessibility logits per cell (Fig.S6). L2 regularization can be optionally applied to the cell-embedding path (with hyperparameter *λ*_1_) or the batch-specific path (with hyperparameter *λ*_2_) to tune the contribution of the batch covariate to the predictions.
- Final sigmoid activation to [0,1] accessibility probability.

The total number of trainable parameters in the model is a function of the number of cells in the dataset. Specifically, the model will have 4513960+33 × n cells number of trainable parameters.

### 5.4 Training approach

We used a binary cross entropy loss and monitored the training area under the receiver operator curve (auROC) after every epoch. We stopped training when the maximum training auROC improved by less than 1e-6 in 50 epochs. This stopping criteria led to training for around 600 epochs for the Buenrostro2018 dataset, 1100 epochs for the 10x multiome PBMC dataset and 1200 epochs for the 10x multiome mouse brain dataset.

We focused on training auROC instead of validation auROC for model selection because we observed that the model continues to improve cell embeddings even after the point where the validation auROC has plateaued (Fig.S14). Since our goal in this application is to learn better representations instead of minimum generalization loss, we focused on the convergence of the training auROC. In addition, at the bottleneck size of 32, there was only a small drop in generalization performance (validation auROC) when the training auROC reaches convergence (0.734 versus 0.742).

We updated model parameters using stochastic gradient descent using the Adam update algorithm. We performed a random search for optimal hyperparameters including: batch size, learning rate, beta1, and beta2 for the Adam optimizer. The best performance was achieved with a batch size of 128, learning rate of 0.01, beta 1 of 0.95, and beta 2 of 0.9995.

We focused on the Buenrostro2018 dataset to select the optimal bottleneck layer size. We trained models with bottleneck sizes of 8, 16, 32, 64 and 128 and observed that bottleneck size 32 gives the best performance (Fig.S13).

### 5.5 Alternative scATAC-seq methods

#### 5.5.1 PCA

We performed PCA with the scikit-learn python package. We evaluated the performance of PCA cell embedding using 10, 20, 30, 40, 50, 60, 70, 80, 90, 100 PCs, with or without the first PC, and reported the model with best performance to compare to scBasset.

#### 5.5.2 cicero

We used Cicero via its R package (Pliner et al., 2018). We ran preprocess cds() function on the binarized peak by cell matrix with method=‘LSI’, followed by reducedDims() function to learn a vector representation for each cell. PCs whose Pearson correlations with sequencing depth*>*0.5 are removed.

#### 5.5.3 cisTopic

We used cisTopic via its R package (Bravo Gonz’
salez-Blas et al., 2019). We ran runCGSModels() function on the binarized peak by cell matrix with a range of topic numbers (2, 5, 10, 20, 30, 40, 50, 60, 80 and 100) for 200 iterations with burn in periods of 120. For comparison with scBasset, we reported the cisTopic models with the best cell embedding performance.

#### 5.5.4 SCALE

We used SCALE via its command line tool (https://github.com/jsxlei/ SCALE) with parameters -x 0.05 and -min peaks 500 to filter low quality peaks and cells to avoid exploding gradients (Xiong et al., 2019). We ran SCALE with a range of latent sizes (10, 16, 32, 64) and found that the default latent size of 10 gives the best cell embedding performance. We also added the –impute option allowing SCALE to estimate denoised accessibility values.

#### 5.5.5 chromVAR

We used ChromVAR via its R package (Schep et al., 2017). We first created a summarized experiment object from the binary peak by cell matrix, followed by addGCBias() using the corresponding genome build. We featurize the sequences into motif space using Jaspar motifs or k-mer space using 6-mers. Next, we computed the deviation z-score matrices for motif and k-mer matches. For each of chromVAR-motif or chromVAR-kmer, we performed PCA on the motif deviation score matrix with 10, 20, 30, 40, 50, 60, 70, 80, 90, 100 PCs and reported the best cell embedding performance to compare to scBasset.

When using chromVAR for TF activity inference, we ran chromVAR motif match using CIS-BP motifs instead of the default Jaspar motifs for a fair comparison with scBasset. Then we computed deviation z-scores as previously described.

### 5.6 Cell embedding evaluation

Adjusted rand index (ARI): We evaluated learned cell embeddings in the Buenrostro2018 dataset by comparing the clustering to the ground-truth cell type labels. We first built a nearest neighbor graph using scanpy with default n neighbors of 15. Then we followed a previous study to tune for a resolution that outputs 10 clusters (Chen et al., 2019). Finally, we compared the clustering outcome to the ground-truth cell type labels using ARI.

Label score: We evaluated the learned cell embeddings using label score for all three datasets. For a given nearest neighbor graph, label score quantifies what percentage of each cell’s neighbors share its same label in a given neighborhood. For each cell embedding method, we computed label score across a neighborhood of 10, 50 and 100. Since the ground-truth cell types for the multiome datasets are unknown, we used cluster identifiers from scRNA-seq Leiden clustering as cell type labels.

Neighbor score: We evaluated the learned cell embeddings using neighbor score for the 10x multiome datasets. For a 10x multiome dataset, we built in-dependent nearest neighbor graphs from the scRNA (using scVI) and scATAC (using the cell embedding method we want to evaluate) and quantified the percentage of each cell’s neighbors that are shared between the two graphs across neighborhoods of size 10, 50 and 100.

### 5.7 Batch correction evaluation

We evaluated scBasset-BC on additional scATAC datasets from mixed PBMC populations from 10x PBMC multiome chemistry (downloaded from https://cf.10xgenomics.com/samples/cell-arc/1.0.0/pbmc_granulocyte_sorted_10k/) and 10x PBMC nextgem chemistry (https://cf.10xgenomics.com/samples/cell-atac/2.0.0/atac_pbmc_10k_nextgem/). We generated a shared atlas of 21,017 peaks from the two datasets by resizing the 10x peak calls from the two datasets to 1000bp and took the intersection. We subsampled 2,000 cells from each dataset and merged them over the shared atlas. We ran scBasset-BC with hyperparameters *λ*_1_=1e-6 and *λ*_2_=0.

### 5.8 Denoising evaluation

To compute a denoised and normalized accessibility across cells for a query peak with scBasset, we ran a forward pass on the input DNA sequence to compute the latent embedding for the peak. Then we generate the normalized accessibility across all cells through dot product of the peak embedding with the weight matrix of the final layer. Notice that since sequencing depth information is entirely captured by the intercept vector of the final layer, we excluded the intercept term so that scBasset generates denoised profiles normalized for sequencing depth.

Our evaluation is based on the hypothesis that effective denoising would improve the correlation between accessibility at genes’ promoters and the genes’ expression in the multiome measurements (Granja et al., 2021; Pliner et al., 2018). For each gene, we computed a gene accessibility score by averaging the accessibility values for peaks at the gene’s promoter (±2kb from TSS). We evaluated denoising performance by computing the Pearson correlation between the gene accessibility score and gene expression (after scVI denoising) across all genes for each individual cell.

Alternatively, we also evaluated scBasset accessibility denoising for consistency between differential expression and differential accessibility. We performed differential gene expression on scVI gene expression for each cell type cluster versus the rest. We also performed differential accessibility analysis on gene accessibility scores for each cell type cluster versus the rest. Then we evaluated performance by computing the Pearson correlation between the gene accessibility score log2FC and gene expression log2FC across all genes for each cell type cluster.

### 5.9 Integration evaluation

In order to evaluate integration performance, we treated the 10x multiome scRNA and scATAC profiles as originated from two independent experiments. We summarize the accessibility profile to a gene level by computing gene accessibility score as described above and integrated the scRNA and scATAC data by embedding them into a shared space using Seurat FindTransferAnchors() and TransferData() functions (Stuart et al., 2019).

In order to quantify the integration performance, we measured a “multiome rank distance” R_c_ between the RNA embedding and the ATAC embedding of each cell *c*. We use R_rna_ to represent the ranking of the Euclidean distance between RNA embedding and ATAC embedding of cell *c* among all neighbors of *c*’s RNA embedding, and R_atac_ to represent the ranking of the same distance among all neighbors of *c*’s ATAC embedding. R_c_ is computed as the average of R_rna_ and R_atac_.

### 5.10 Motif injection

We performed motif injection on scBasset to compute a TF activity score for each TF for each cell. Specifically, we first generated 1000 genomic background sequences by performing dinucleotide shuffling of 1000 randomly sampled peaks from the atlas using fasta ushuffle (Jiang et al., 2008). For each TF in the motif database, we sampled a motif sequence from the position weight matrix (PWM) and inserted into the center of each of the genomic background sequences. We ran forward passes through the model for both the motif-injected sequences and background sequences to predict normalized accessibility across all cells. We took the difference in predicted accessibility between the motif-injected sequences and background sequences as the motif influence for each sequence. We averaged this influence score across all 1000 sequences for each cell to generate a cell level prediction of raw TF activity. Finally, we z-score normalized the raw TF activities to generate the final TF activity predictions across all cells.

We used CIS-BP 1.0 single species DNA database motifs downloaded from https://meme-suite.org/meme/db/motifs for our motif analysis (Weirauch et al., 2014).

### 5.11 *In silico* saturation mutagenesis

We performed *in silico* saturation mutagenesis (ISM) to compute the importance scores of all single nucleotides on a sequence of interest. For each position, we ran three scBasset forward passes, each time mutating the reference nucleotide to an alternative. For each mutation, we compared the accessibility prediction to the prediction with the reference nucleotide to compute the change in accessibility for each cell. We normalized the ISM scores for the four nucleotides at each position such that they sum to zero. We then took the normalized ISM score at the reference nucleotide as the importance score for that position.

## 6 Code Availability

Code for training and using scBasset model can be found at: https://github.com/calico/scBasset.

## 7 Acknowledgements

We thank Vikram Agarwal, Jacob Kimmel and Majed Mohamed for feedback on the manuscript. We thank Sarah Spock for feedback on the code. We also thank Nick Bernstein and Ashlesha Odak for helpful discussions.

## 8 Author Contributions

D.R.K. conceived the project. H.Y. and D.R.K. developed the model. H.Y. performed the analysis. H.Y. and D.R.K prepared the manuscript.

## 10 Supplementary Figures

**Figure S1:**
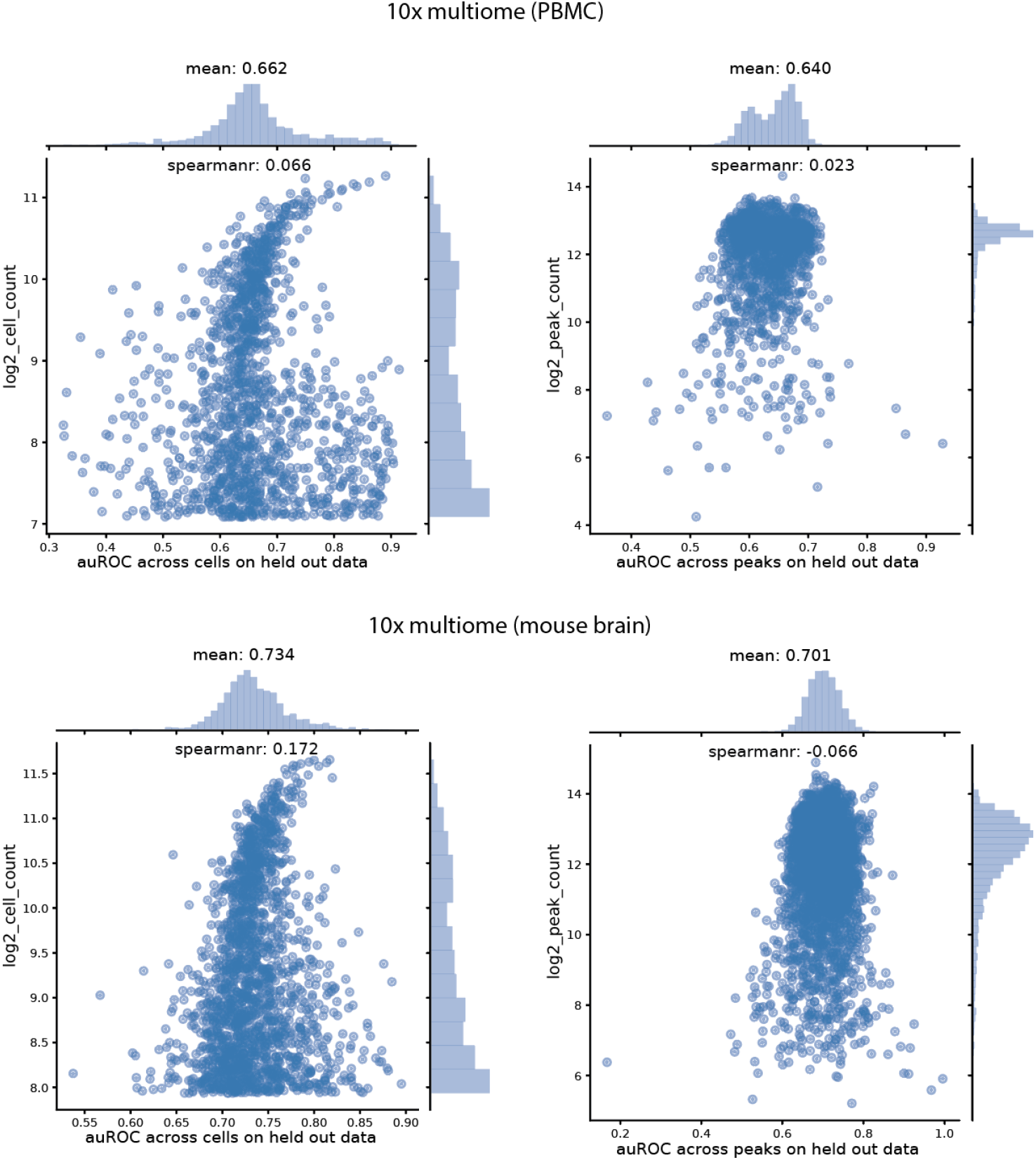
auROC on held-out peaks for 10x multiome PBMC and mouse brain datasets. Top, scBasset prediction performance on held-out peaks evaluated by auROC per peak (left) and by auROC per cell (right) for 10x multiome PBMC dataset. Bottom, scBasset prediction performance on held-out peaks evaluated by auROC per peak (left) and by auROC per cell (right) for 10x multiome mouse brain dataset.

**Figure S2:**
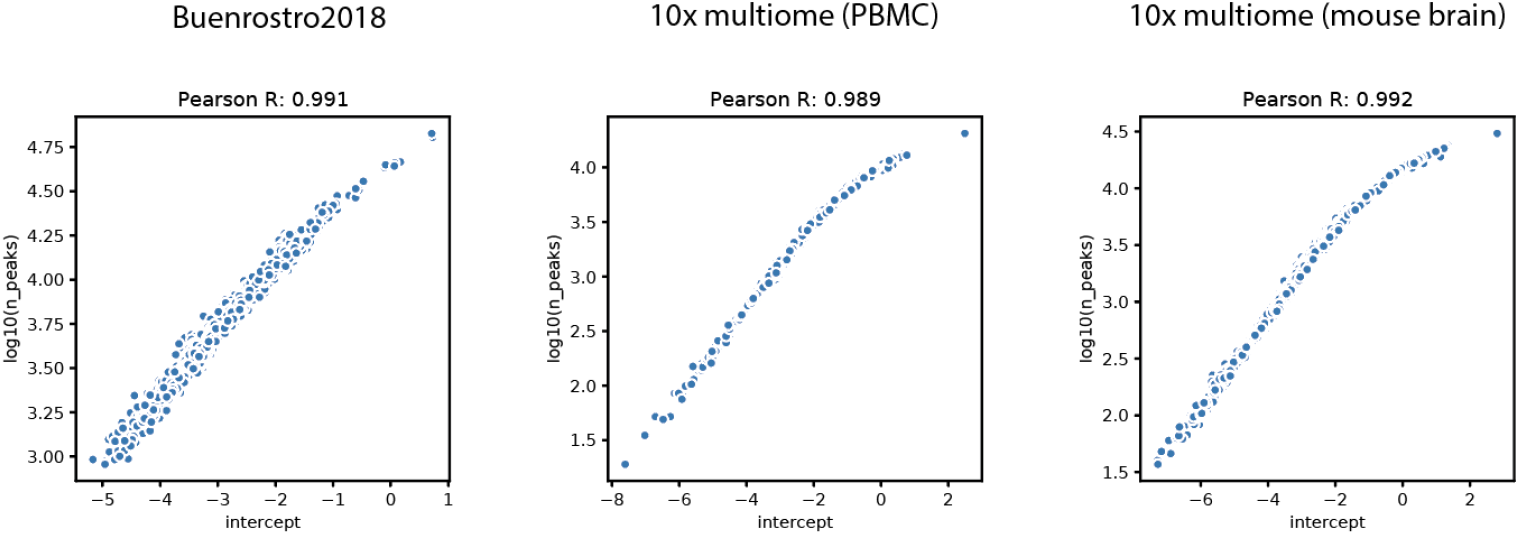
Correlations of final layer intercepts with sequencing depth (log10 UMI) for Buenrostro2018, 10x multiome PBMC and 10x multiome mouse brain datasets (from left to right).

**Figure S3:**
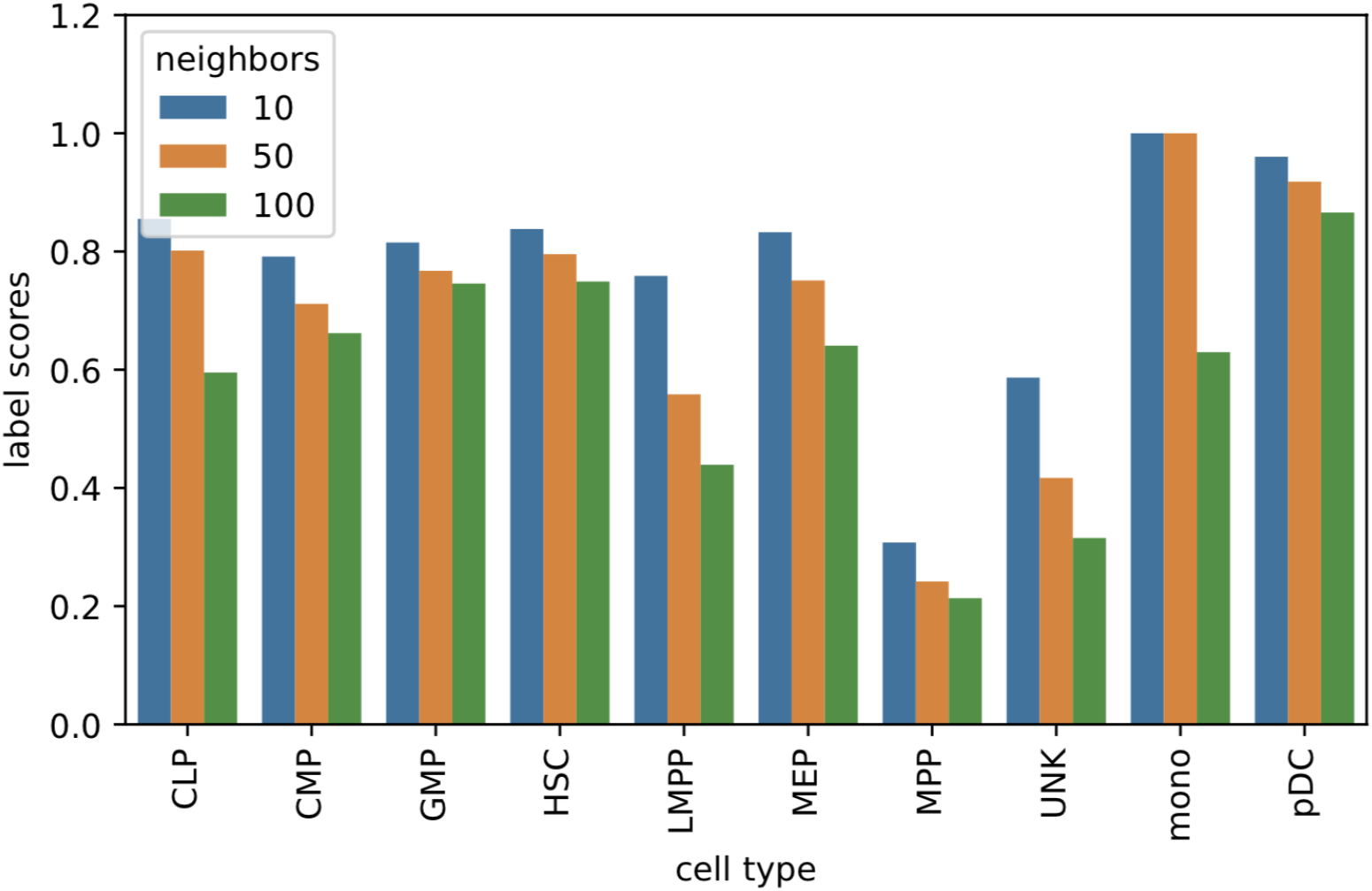
scBasset cell embedding performance as evaluated by label scores for each cell type with a neighborhood of 10, 50 and 100.

**Figure S4:**
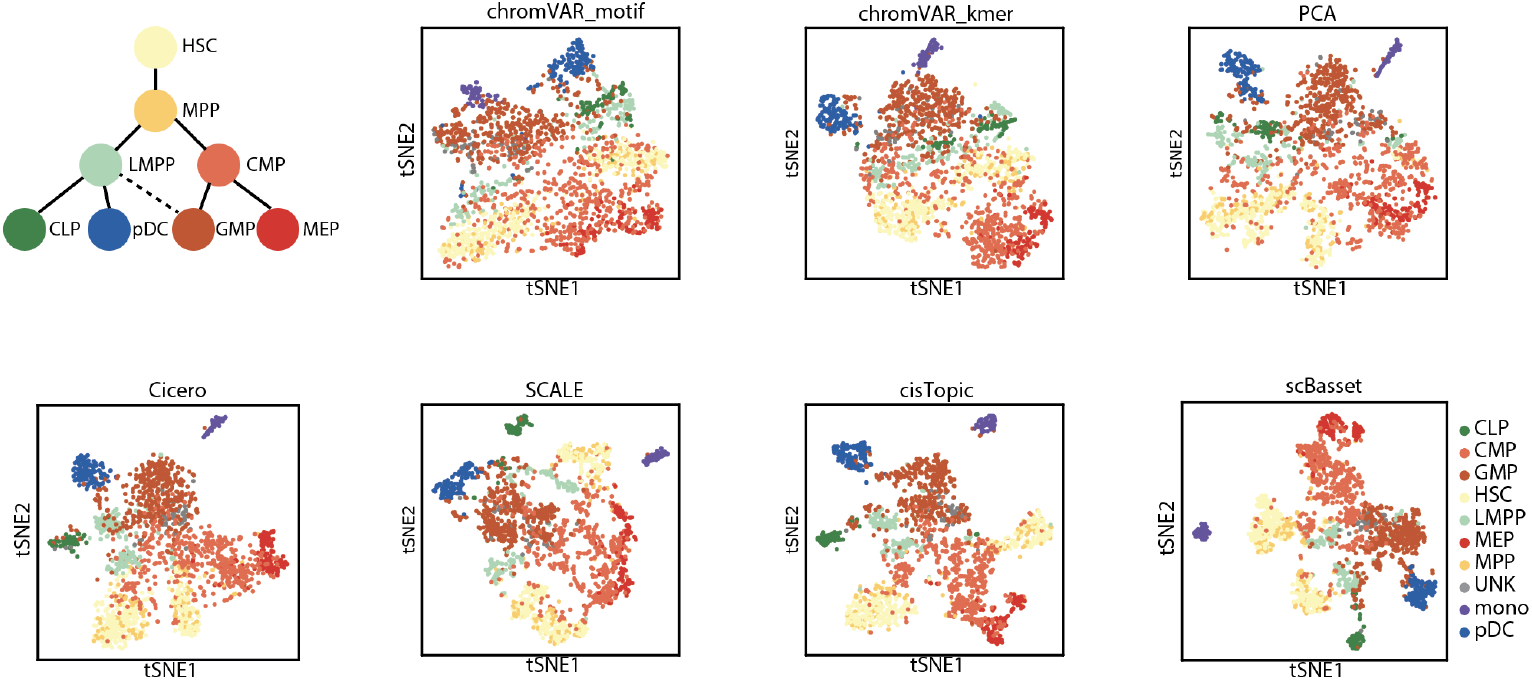
t-SNE visualization of different cell embedding methods, including: chromVAR motif, chromVAR kmer (k=6), PCA, cicero (LSI), SCALE, cisTopic and scBasset.

**Figure S5:**
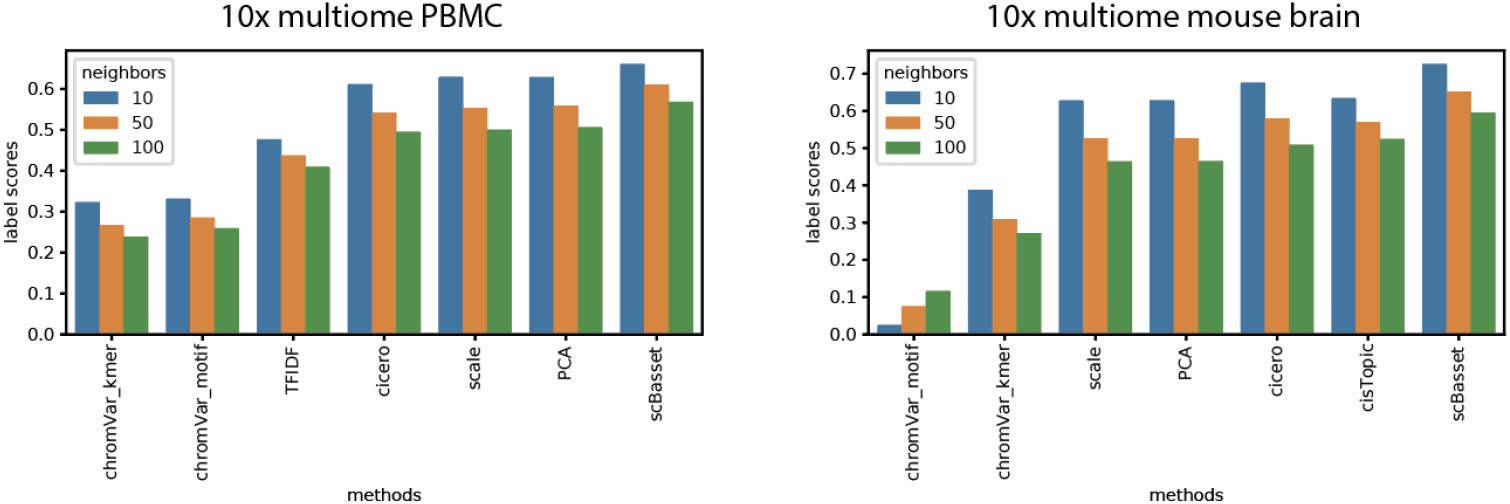
Performance comparison of different cell embedding methods as evaluated by label scores for 10x multiome PBMC (left) and mouse brain (right) datasets.

**Figure S6:**
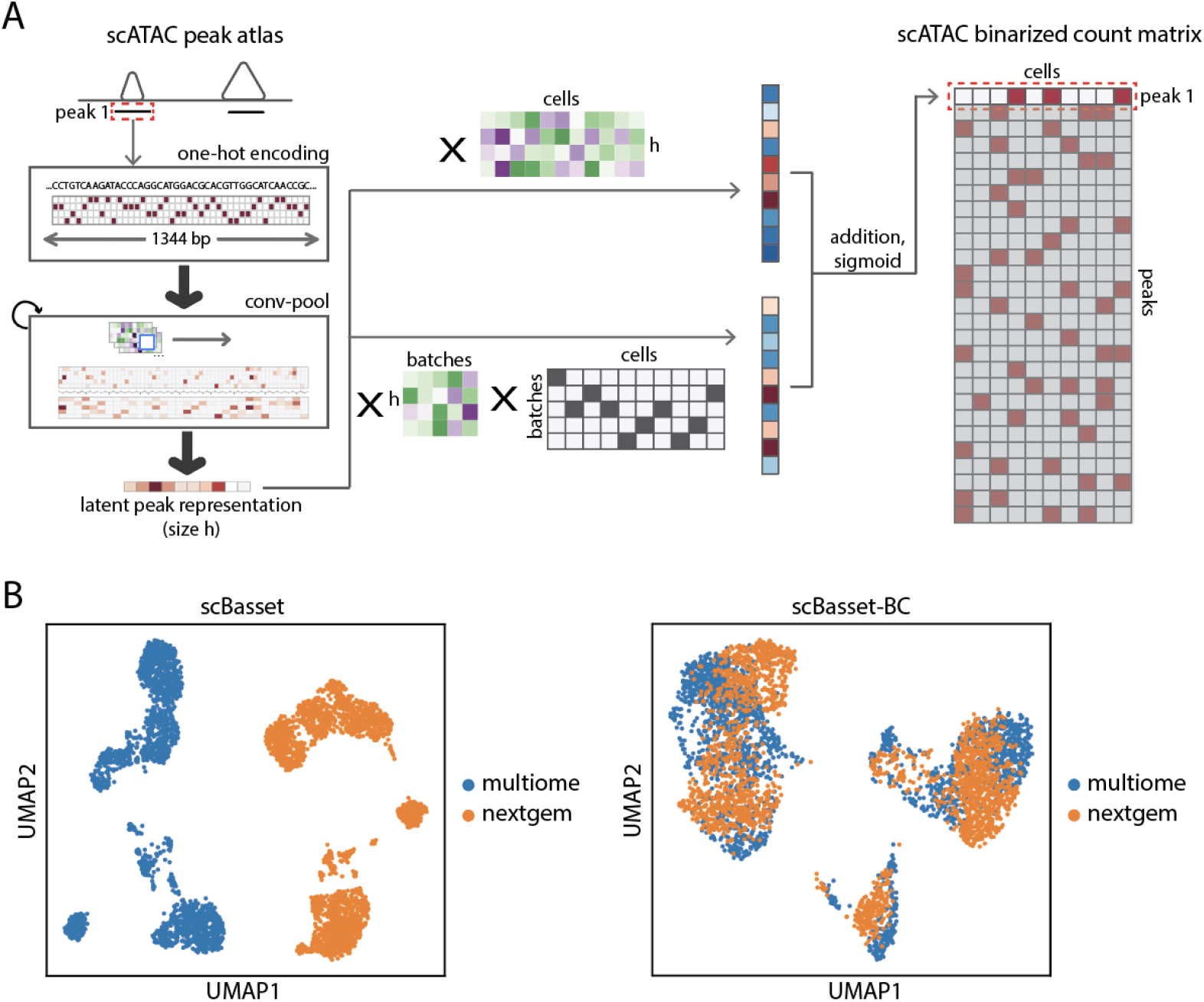
A, Model architecture of scBasset-BC. B) UMAP embeddings of mixed PBMC populations from 10x multiome scATAC and 10x nextgem scATAC chemistries. Left figure shows the embeddings learned by scBasset model. Right figure shows the embeddings learned by scBasset-BC model, where batch is encoded as a covariate.

**Figure S7:**
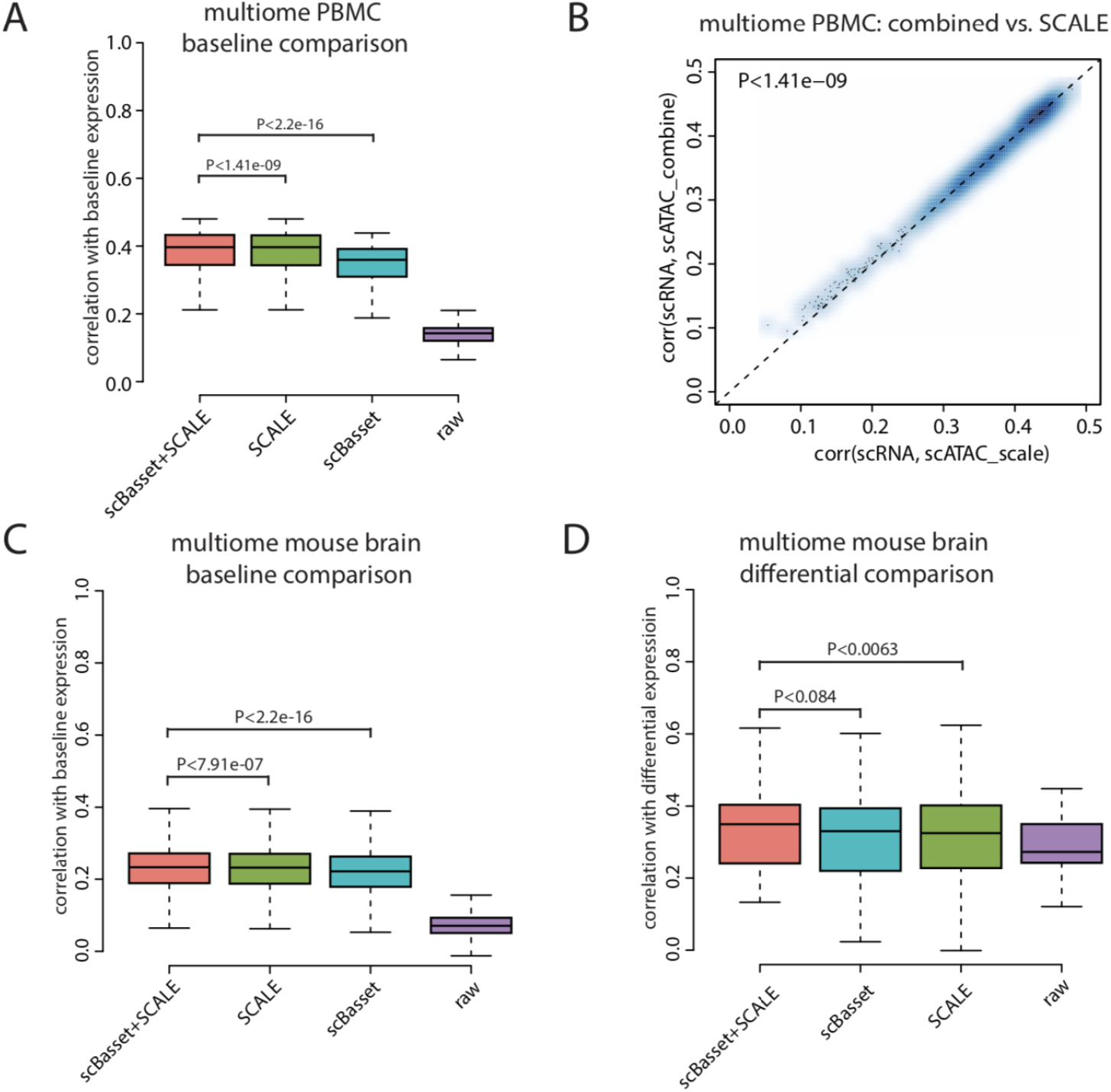
Additional denoising results for 10x multiome datasets. A) Comparison of denoising performance on the multiome PBMC dataset between raw data, scBasset, SCALE, and scBasset+SCALE combined, evaluated by correlation between baseline gene accessibility score and baseline gene expression. B) A scatterplot showing a closer look at the performance comparison between scBasset+SCALE (y-axis) versus SCALE on multiome PBMC dataset, evaluated by correlation between baseline gene accessibility score and baseline gene expression. C) Comparison of denoising performance on the multiome mouse brain dataset between raw data, scBasset, SCALE, and scBasset+SCALE combined, evaluated by correlation between baseline gene accessibility score and baseline gene expression. D) Comparison of denoising performance on multiome mouse brain dataset between raw data, scBasset, SCALE, and scBasset+SCALE combine, evaluated by consistency in differential expression log2FC and differential accessibility log2FC. We performed Wilcoxon signed rank tests for performance comparisons.

**Figure S8:**
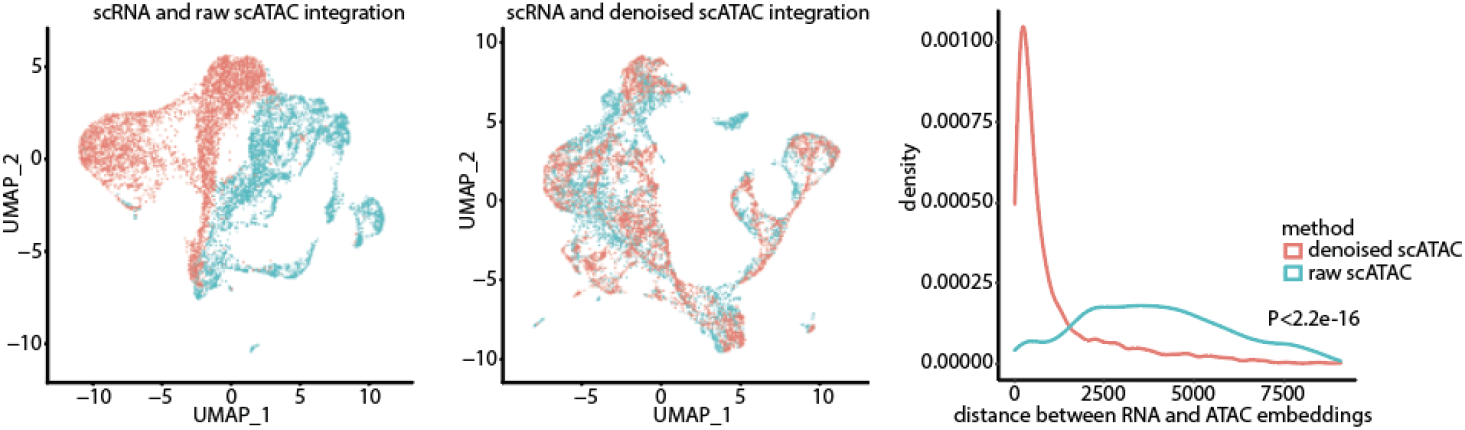
Integration results for the 10x multiome mouse brain dataset. Left, RNA (blue) and raw ATAC (red) profile embeddings after integration. Middle, RNA (blue) and denoised ATAC (red) profile embeddings after integration. Right, distribution of the relative distances (Methods) between each cell’s RNA and ATAC embeddings after integration when integrating with raw ATAC profiles (blue) or denoised ATAC profiles (red). We performed Wilcoxon signed rank test for performance comparison.

**Figure S9:**
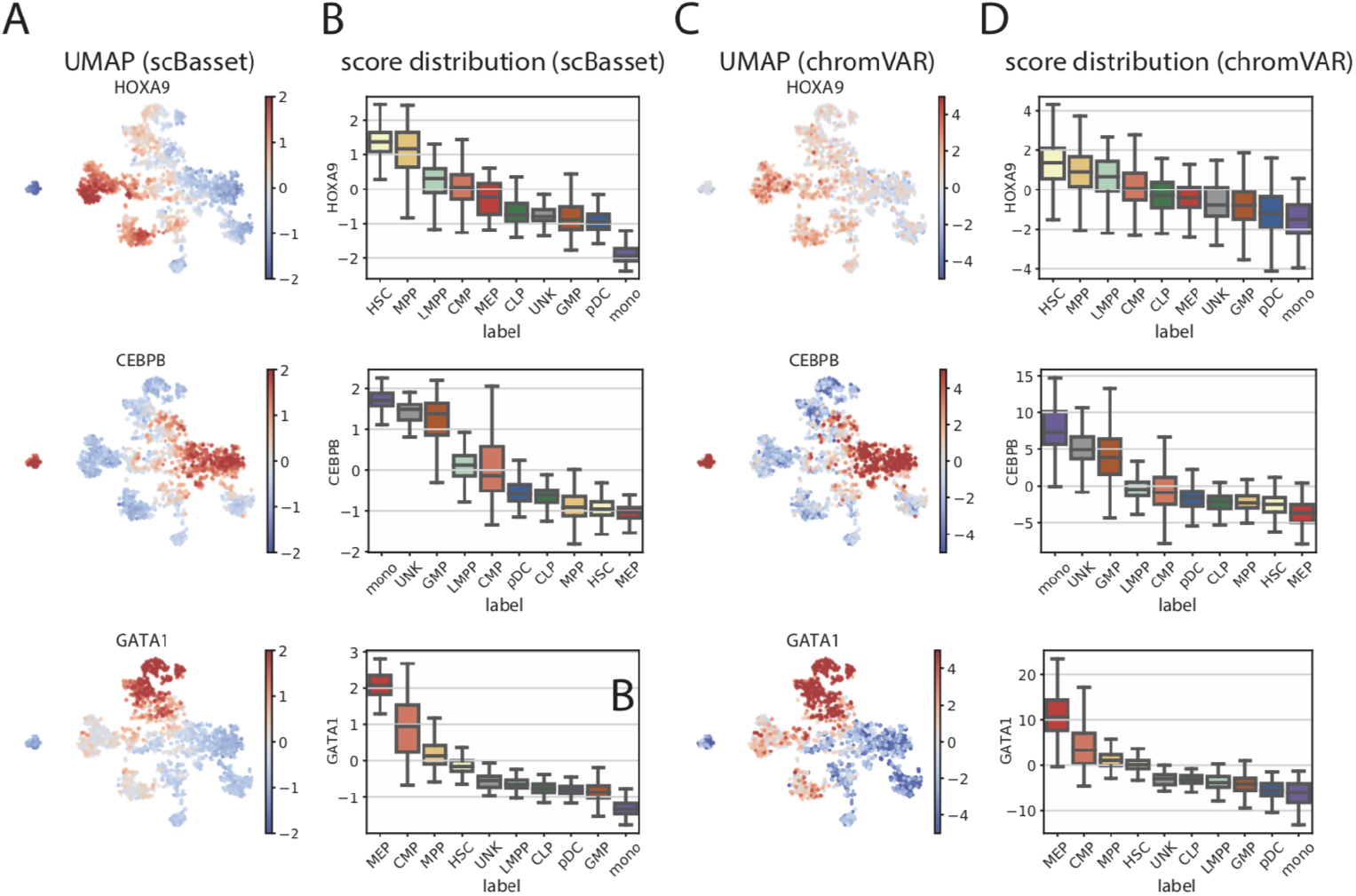
Motif activity inference using scBasset and chromVAR on the Buenrostro 2018 dataset for known regulators. A) UMAPs showing scBasset-predicted TF activity. B) Boxplots showing scBasset-predicted TF activity by cell type. C) UMAPs showing chromVAR-predicted TF activity. B) Boxplots showing chromVAR-predicted TF activity per cell type.

**Figure S10:**
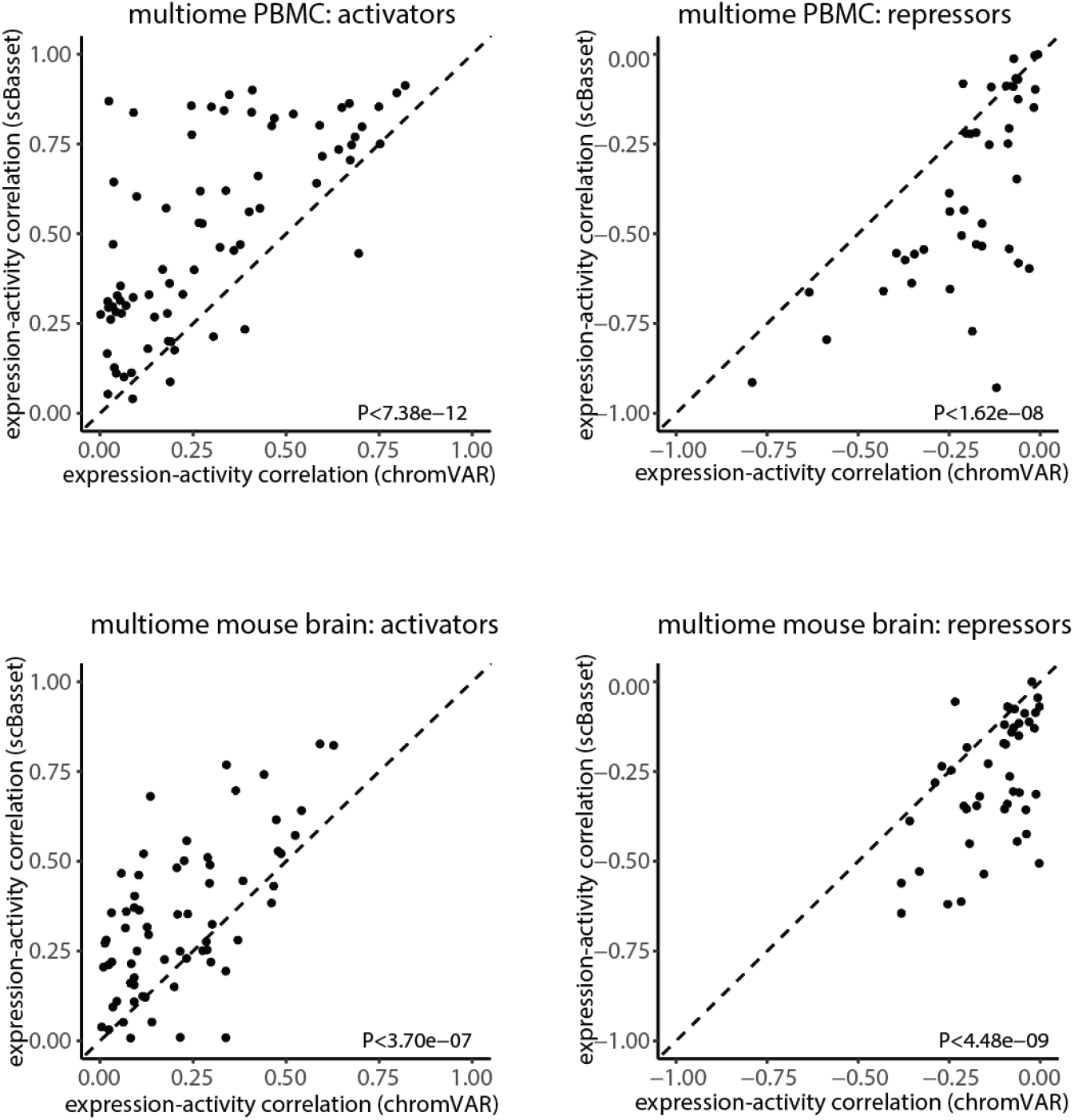
TF expression and TF activity correlation for the 10x mulitome datasets. Scatterplots of correlations between chromVAR-inferred activity and expression (x-axis) versus correlations of scBasset-inferred TF activity and expression (y-axis) for activating TFs (left) and repressive TFs (right) in the 10x multiome PBMC (top) and 10x multiome mouse brain (bottom). Activating TFs are TFs which both scBasset and chromVAR agree on a positive correlation between TF expression and activity. Repressive TFs are TFs which both scBasset and chromVAR agree on a negative correlation between TF expression and activity.

**Figure S11:**
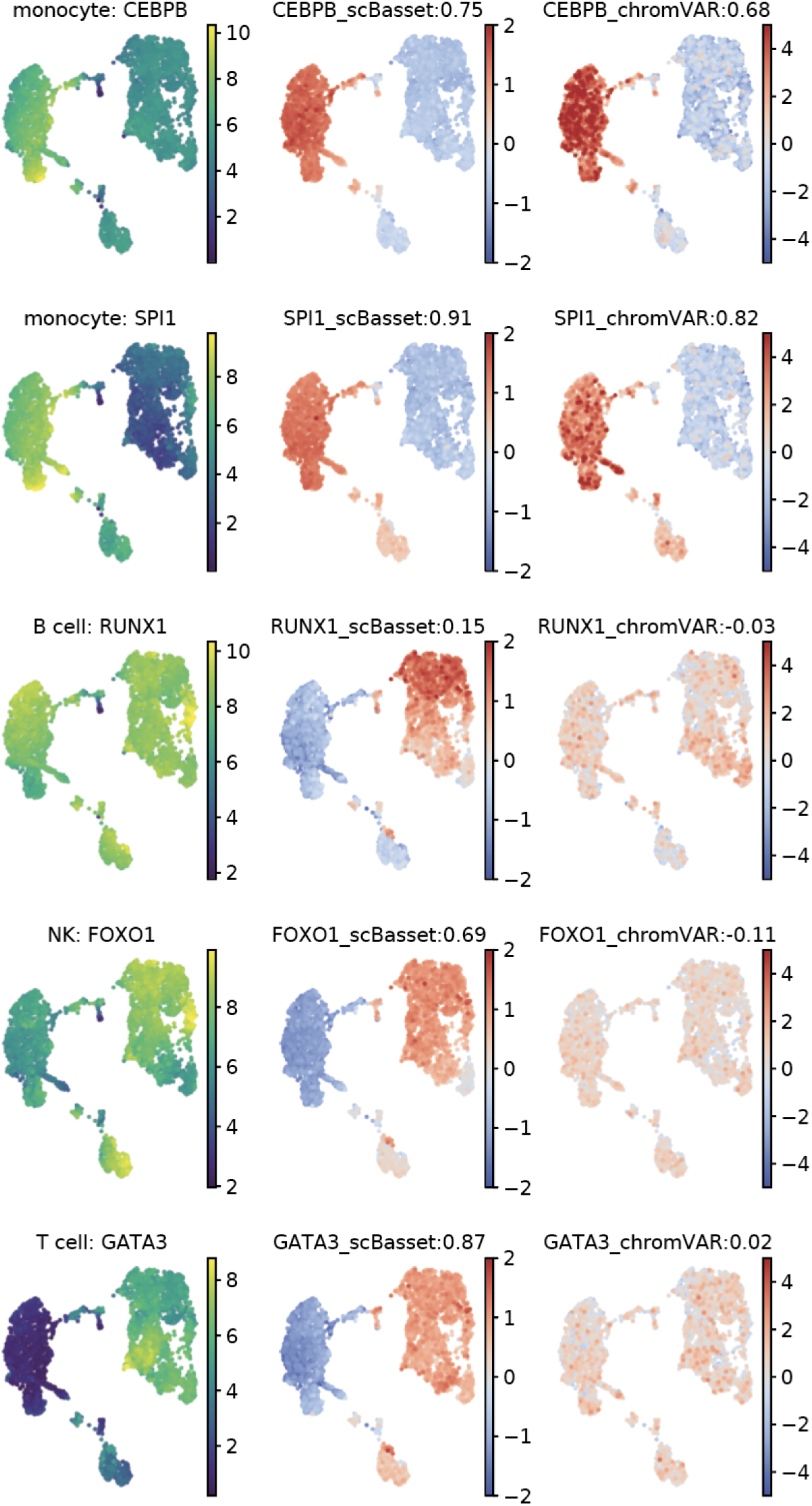
Motif activity inference using scBasset and chromVAR on the 10x multiome PBMC data. UMAP visualization of TF expression (left), scBasset TF activity (middle), and chromVAR TF activity (right) for additional known PBMC regulators. Pearson correlation between inferred TF activity and expression are shown in the titles.

**Figure S12:**
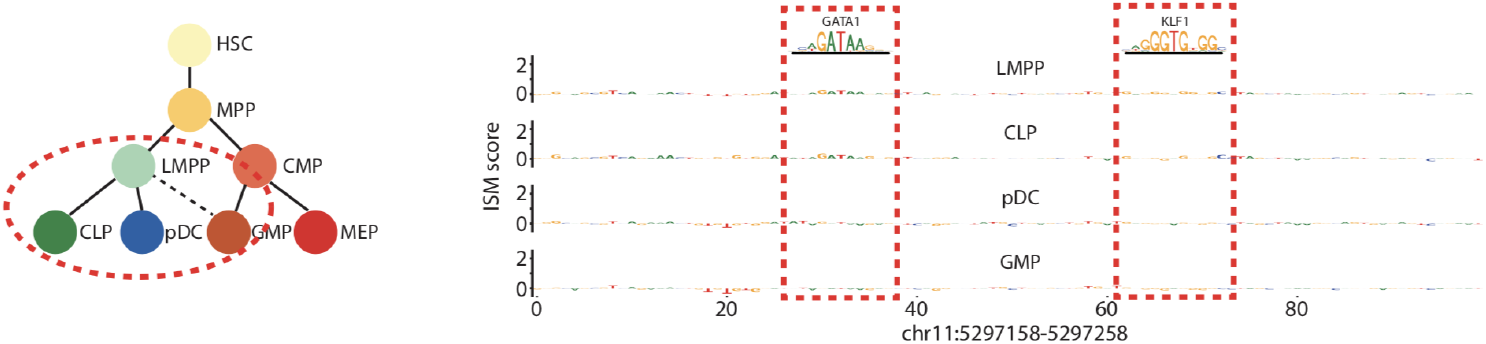
ISM scores for *β*-globin enhancer at chr11:5297158-5297258 for cells in LMPP, CLP, pDC and GMP cell types. Sequences that match GATA1 and KLF1 motifs are highlighted in red boxes.

**Figure S13:**
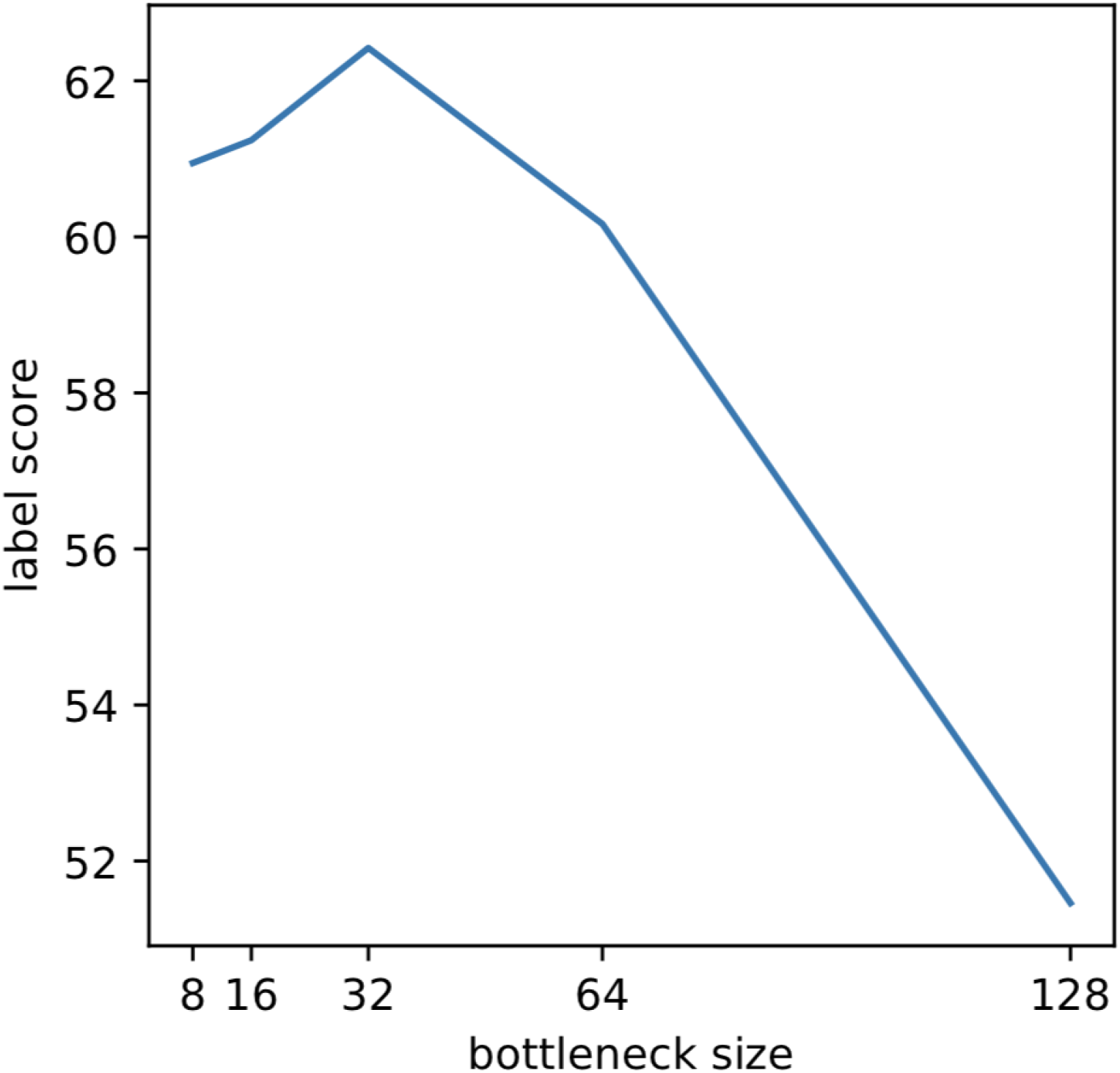
Label scores as a function of scBasset bottleneck layer size in Buenrostro2018 dataset.

**Figure S14:**
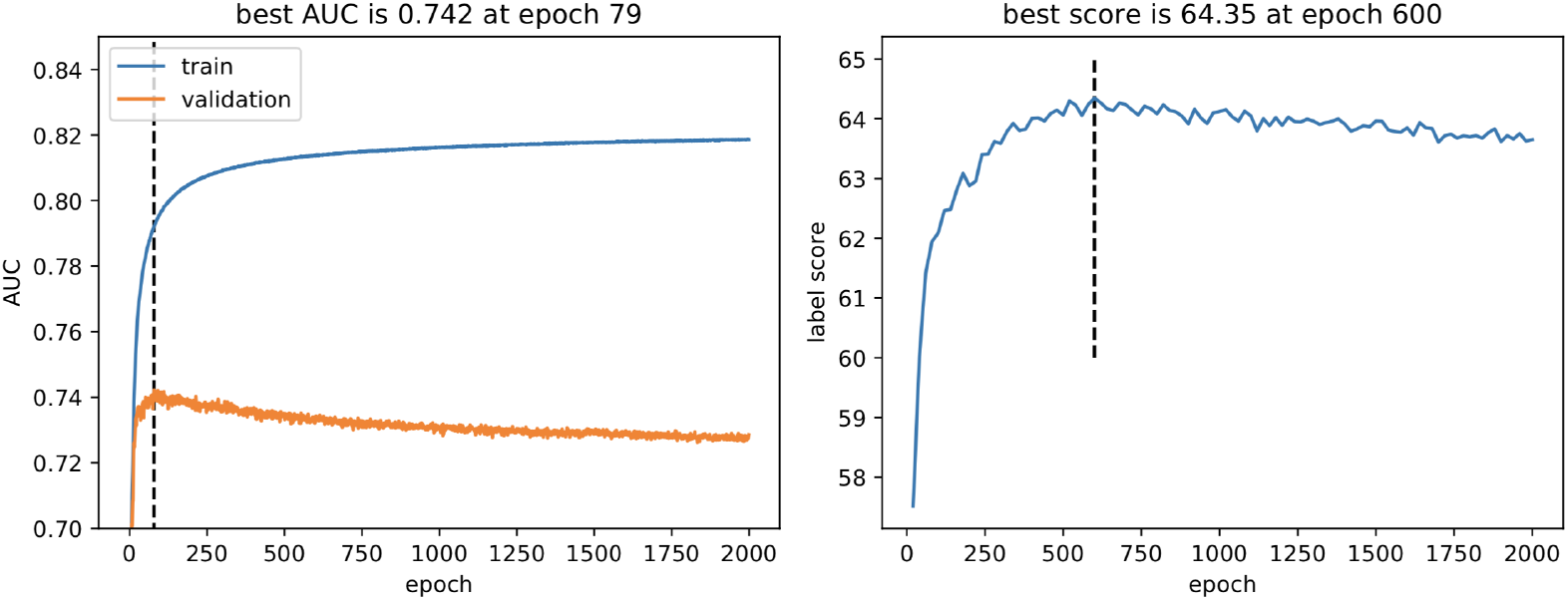
Left, training (blue) and validation auROCs (red) per epoch for the Buenrostro2018 dataset. Right, label scores per epoch.

